# Comparison of two different finite element modeling pipelines for virtual mechanical testing of the distal third metacarpal bone in Thoroughbred racehorses

**DOI:** 10.1101/2025.10.29.685454

**Authors:** Soroush Irandoust, Fatemeh Malekipour, R. Christopher Whitton, Peter Muir, Peter Vee-Sin Lee, Corinne R. Henak

## Abstract

Condylar stress fracture of the third metacarpal bone (MC3) in Thoroughbred racehorses is a common catastrophic injury and identification of horses at heightened risk remains subjective. Standing computed tomography (sCT) is a practical screening tool that is sensitive to fatigue-induced structural changes. Data from sCT also allows for patient-specific finite element analysis (FEA) of the distal MC3 and prediction of subchondral bone strain, as a potential objective classifier of racehorses at heightened risk. The goal of this study was to compare two independently developed sCT-based subject-specific FEA pipelines for virtual mechanical testing of the distal MC3. One pipeline models the full 3D distal MC3 (UWMSN), while the other uses a simpler approach by using single sCT slices (UMELB). Four (n=4) MC3 condyles from four Thoroughbred racehorses were selected for the study. Models were generated using both pipelines and the predicted subchondral bone strain was compared. UMELB predicted smaller subchondral strain compared to UWMSN, likely due to more limited modes of deformation. Although the UWMSN pipeline can identify elevated subchondral strain in horses with high fatigue damage, it is more labor intensive and computationally expensive. With further tuning and validation, the UMELB pipeline could be used as a simpler and faster approach for prediction of subchondral strain in the distal MC3.

## 1. Introduction

Condylar stress fracture of the metacarpophalangeal (MC3, fetlock) joint is a common catastrophic injury in Thoroughbred racehorses worldwide and a major cause of euthanasia [1,2] and jockey falls [3]. Thoroughbreds with a complete displaced condylar stress fracture have a significantly decreased prognosis for racing after surgery [4–6], hence, prevention of these injuries and associated horse death is an important focus.

Clinical interpretation of fatigue-induced structural changes in the subchondral bone (SCB) and assessment of the associated condylar fracture risk is currently challenging [7] and an objective classifier is lacking. Standing computed tomography (sCT) imaging is a practical noninvasive racehorse screening method [8,9] and provides the opportunity to build patient-specific finite element (FE) models for virtual mechanical testing. Such models have the potential to improve identification of MC3 bones at heightened risk of fracture.

Most recently, our groups have independently developed FE analysis (FEA) pipelines for virtual mechanical testing of the distal MC3 bone [10,11] using different software packages for image segmentation and FEA. One requires three-dimensional (3D) segmentation of the distal MC3 and building a full 3D FE model (UWMSN), while the other requires segmentation of only slices of the sCT image set and building a slice-based FE model (UMELB). The UMELB pipeline is faster to build and is computationally cheaper to run but does not fully capture the heterogeneity of bone mineral density in the distal MC3 bone.

To enable consistent classification of horses at risk of condylar stress fracture, different existing pipelines need to be benchmarked against each other. This is necessary to improve the reproducibility of the resulting clinical predictions, and accurate modeling has been a recent focus of various regulatory bodies [12,13]. Benchmarking pipelines against each other is rarely done, and a challenge to do well. For example, a recent cohort modeling contact mechanics in the human knee found that many of the steps in developing pipelines result in divergent predictions [14]. To our knowledge, this has not been attempted for modeling pipelines of the equine fetlock.

Therefore, the goal of this study was to compare the predicted strain in the distal MC3 bone between UWMSN and UMELB pipelines.

## 2. Methods

### 2.1. Specimen selection

A library of four MC3 condyles (n=4) from four thoracic limb specimens were used for this study. Specimens were collected from Thoroughbred racehorses that were euthanatized because of catastrophic racetrack injury. Age, sex, and training history of the specimens were not available.

### 2.2. sCT acquisition and calibration

The sCT scans of the frozen limbs were acquired using an Asto CT (Middleton, WI, USA) Equina^®^ scanner at exposure of 160 kVp and 8 mA, with 0.55 mm slice thickness, and 0.7324 mm pixel spacing in each slice. An electron density phantom (model 062M, CIRS Inc, Arlington, VA) with 4 calcium hydroxyapatite (HA) plugs with densities of 200, 800, 1250, and 1750 mgHA/cm^3^ was scanned with the same scanner at the same exposure settings to find the equivalent radiological or CT density, *ρ*_*CT*_ (*mg_HA_*/*cm*^3^):

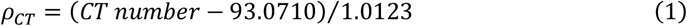

A separate scan was made with a different set of HA plugs with densities of 200, 800, 1250, and 1500 mgHA/cm^3^ with corresponding physical densities of 1.16, 1.53, 1.82, and 1.99 g/cm^3^, to find the physical density, *ρ*(*g*/*cm*^3^):

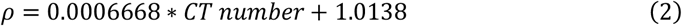

### 2.3. Density-Modulus relationship

Mechanical properties of bone have a strong correlation with its density [15], and this relationship can be used to predict heterogenous bone mechanical properties. First, CT density from equation (1) was converted to ash density using the data from Knowles *et al*. [16]:

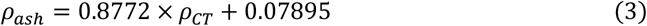

In equation (3) *ρ*_*ash*_ and *ρ*_*CT*_ are expressed in *g*/*cm*^3^. The Young’s modulus was then calculated using the experimental data collected from horse limbs by Moshage *et al*. [17]:

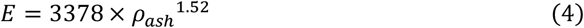

In equation (4) *E* is expressed in *MPa* and *ρ*_*ash*_ in *g*/*cm*^3^.

In addition to the density-modulus relationship in equation (4), the UMELB pipeline used an additional relationship which was specifically developed from compressive mechanical testing of equine MC3 SCB [18]:

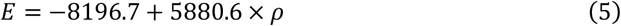

In equation (5) *E* is expressed in *MPa* and *ρ* (calculated from equation (2)) in *g*/*cm*^3^. Equation (5) predicts smaller Young’s moduli than equation (4) (**Figure S1**).

### 2.4. Model preparation: Image segmentation and finite element analysis

The two teams involved in the study started with the same sCT image sets and used the same density-modulus relationship (explained in sections 2.2 and 2.3 above) but developed different pipelines independently using different software packages for image segmentation and FEA (UWMSN [10] and UMELB [11]).

#### 2.4.1. UWMSN pipeline

This pipeline was explained in detail previously [10] and is summarized here. The distal MC3 was segmented using Mimics (v.26, Materialise, Belgium) (**Figure 1**). The external surface of the bone was detected semi-automatically for all individual sCT image slices sequentially and the outer layer of pixels was eroded from the periosteal surface in the end to make sure it was tight on the cortex before smoothing and importing it into 3-matic (v.18, Materialise, Belgium) for further processing. An anatomical coordinate system was established by first finding the long axis of the bone and constructing the transverse, sagittal, and then frontal planes in the same order. The distal 2.5 inches of the MC3 was isolated and a homogenous triangular surface mesh with 0.25 mm edge length was generated.

**Figure 1.**
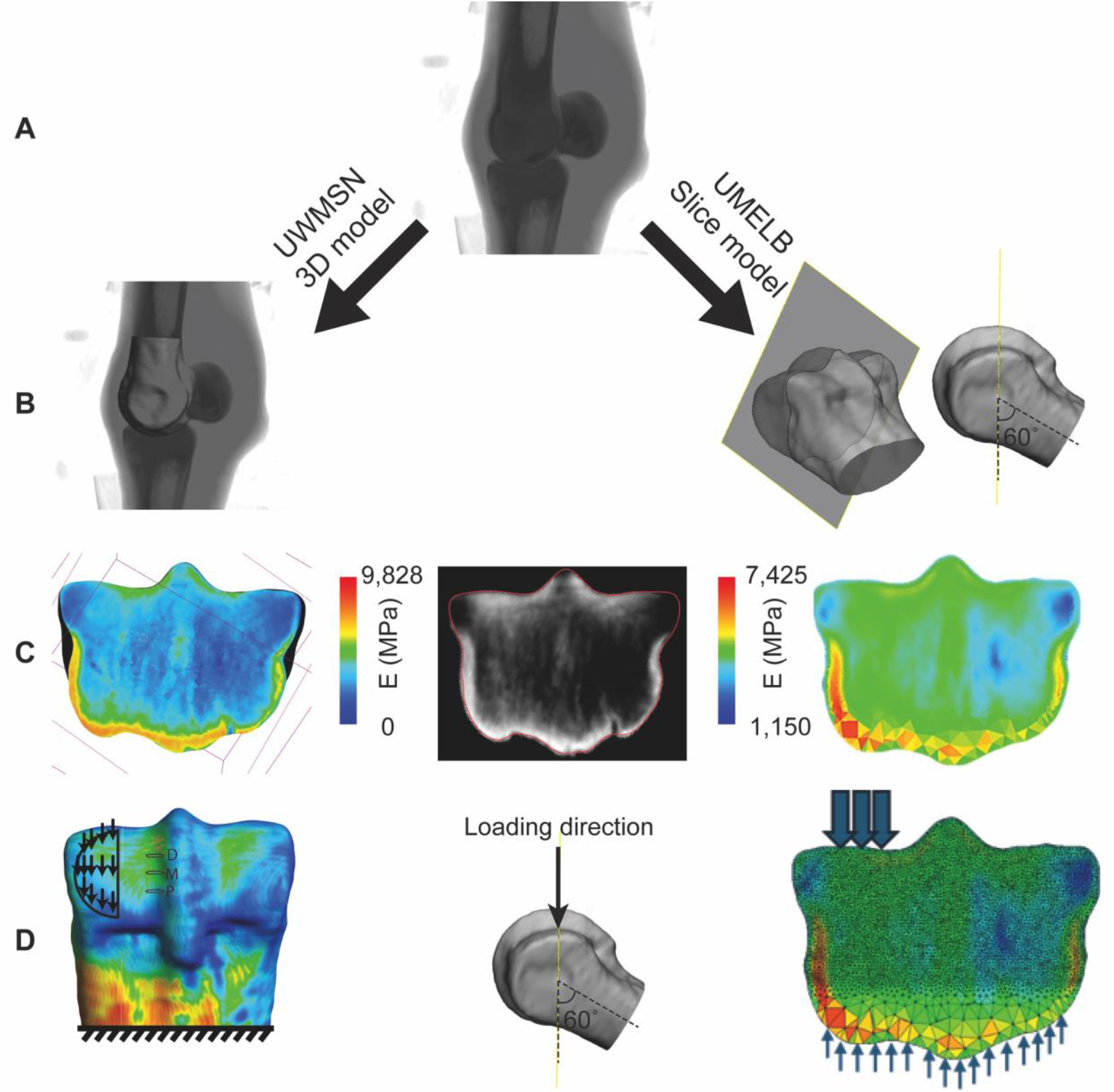
**A)** Standing CT (sCT) image sets were used for the UWMSN (left) and UMELB (right) finite element (FE) modeling pipelines. **B)** The entire distal MC3, and a single oblique slice were segmented in the UWMSN and UMELB pipelines, respectively. **C)** Element-wise heterogenous HA density was captured from the sCT image set and the Young’s modulus was assigned to individual elements. **D)** Loading and boundary conditions were applied. The palmar (P), middle (M), and dorsal (D) parasagittal groove (PSG) regions for which the first principal strain was compared between the two pipelines are shown in the UWMSN pipeline.

FE models were created in FEBio Studio (v.2.3) with 0.25 mm edge length linear tetrahedral elements in the subchondral sclerotic region and 1 mm edge length linear tetrahedral elements elsewhere. A Poisson’s ratio of 0.3 and sCT-based Young’s modulus were assigned to individual elements using equations (1), (3), and (4).

The proximal end of the model was fully constrained and a 30 MPa load was applied to the palmar surface of the medial condyle at 60 and 30 degrees with respect to the frontal and transverse planes, respectively (**Figure 1**). This pipeline requires 5-6 hours for 3D segmentation of the distal MC3 and the subchondral sclerotic region, two hours of FE model preparation, and around five hours of runtime on a high throughput computing cluster [19] with at least 240 GB of RAM.

#### 2.4.2. UMELB pipeline

FE models of three discrete slices were generated using a previously described method [11]. All image processing and segmentation, and FE mesh generation was performed in Simpleware (v Q-2020, Synopsys, USA). A single oblique slice was extracted at 60 and 30 degrees with respect to the frontal and transverse planes, respectively, parallel to the applied load. The external surface of the bone was detected semi-automatically for all individual sCT image slices sequentially and one pixel was removed from the external surface in the end to make sure it was not affected by partial volume effects in CT images. This slice was extruded to 20 mm thickness to generate a 3D volume. The mesh volume was generated in Simpleware for all segmented regions using linear tetrahedral elements (C3D4). A mesh convergence study was performed for one model, and the final element size was set to 0.6 mm within the 20 mm superficial subchondral bone (SCB) of MC3, and 1 mm for the remaining MC3 volume. In addition, the FE results were evaluated using linear versus second-order modified tetrahedral elements (C3D10M) for MC3, and differences remained within 1.5% between element types. The meshed volume was then exported to ABAQUS (v6.14-2, Simulia, USA) for FE model preparation. Articular cartilage elements were later converted to hybrid elements in ABAQUS.

To replicate in-situ boundary conditions, the bone was constrained proximopalmarly and distodorsally, with cartilage margins left free to allow natural lateral expansion. The proximal end of MC3 was fully fixed. To ensure this boundary did not artificially affect SCB stresses, the model was extended proximally by 50 mm to simulate a continuous shaft. Sensitivity analyses were performed with this extended model, and by varying SCB modulus from 500 to 50,000 MPa (0.1× to 10× the average). Stress and strain within the superficial 20 mm SCB changed by no more than 5.5%. Therefore, in all remaining slice-based models, only the proximal face was fully constrained.

Surface pressure of 30 MPa was applied to the joint surface over an area that corresponded to the loaded surface in the UWMSN pipeline over the same lateromedial distance and extruded to the entire 20 mm thickness. sCT-based Young’s modulus was assigned to individual elements using equations (2) and (5). We assumed a Poisson’s ratio of 0.3 [20]. The segmentation and model preparation required minimal manual input, as the slice-based segmentation workflow was largely automated using custom Python scripts within Simpleware’s scripting environment (Synopsys, 2023). Only brief visual verification (∼10 minutes) was needed per slice. Subsequent steps, including extrusion, mesh generation, and refinement, were also scripted, allowing full model generation to be completed within minutes after setup. On average, simulations required 1 hour of wall-clock time on a computing cluster with at least 128 GB of RAM.

The process was repeated for two additional slices per horse, taken in the same oblique orientation as the original slice, offset by 10 voxels (i.e., 7 mm) dorsally and palmary. This resulted in three models per specimen, palmar, middle, and dorsal, to predict SCB strain across these regions (Figure 1). The aim of using three slices was to assess the sensitivity of the results to slice location, particularly regarding the position of peak strain and the relative differences between horses.

### 2.5. Comparison of predicted strain levels in the PSG

Contour plots of the first principal strain for all horses were compared between the two pipelines qualitatively. For quantitative comparison, the PSG regions were selected in the UMELB pipeline first and saved as STL files, and the same region was selected in the UWMSN pipeline. Next, the first principal strain was extracted from these regions and the mean, maximum, and 90^th^ percentile values were compared between the two pipelines. Due to the small sample size, no statistical analysis was performed for comparison of the two pipelines.

## 3. Results

Both modeling pipelines predicted elevated first principal strain (most tensile strain) in the SCB underlying the PSG of the loaded condyle, with the UMELB predicting smaller strain values compared to the UWMSN (**Figure 2**) when using the same density-elastic modulus relationship. UWMSN predicted dorsal propagation of the strain elevation in the grooves, unlike UMELB due to the artificial boundary conditions in this pipeline. Additionally, artificially elevated strain was predicted by UMELB focally on the lateral and medial aspects of the dorsal surface.

**Figure 2.**
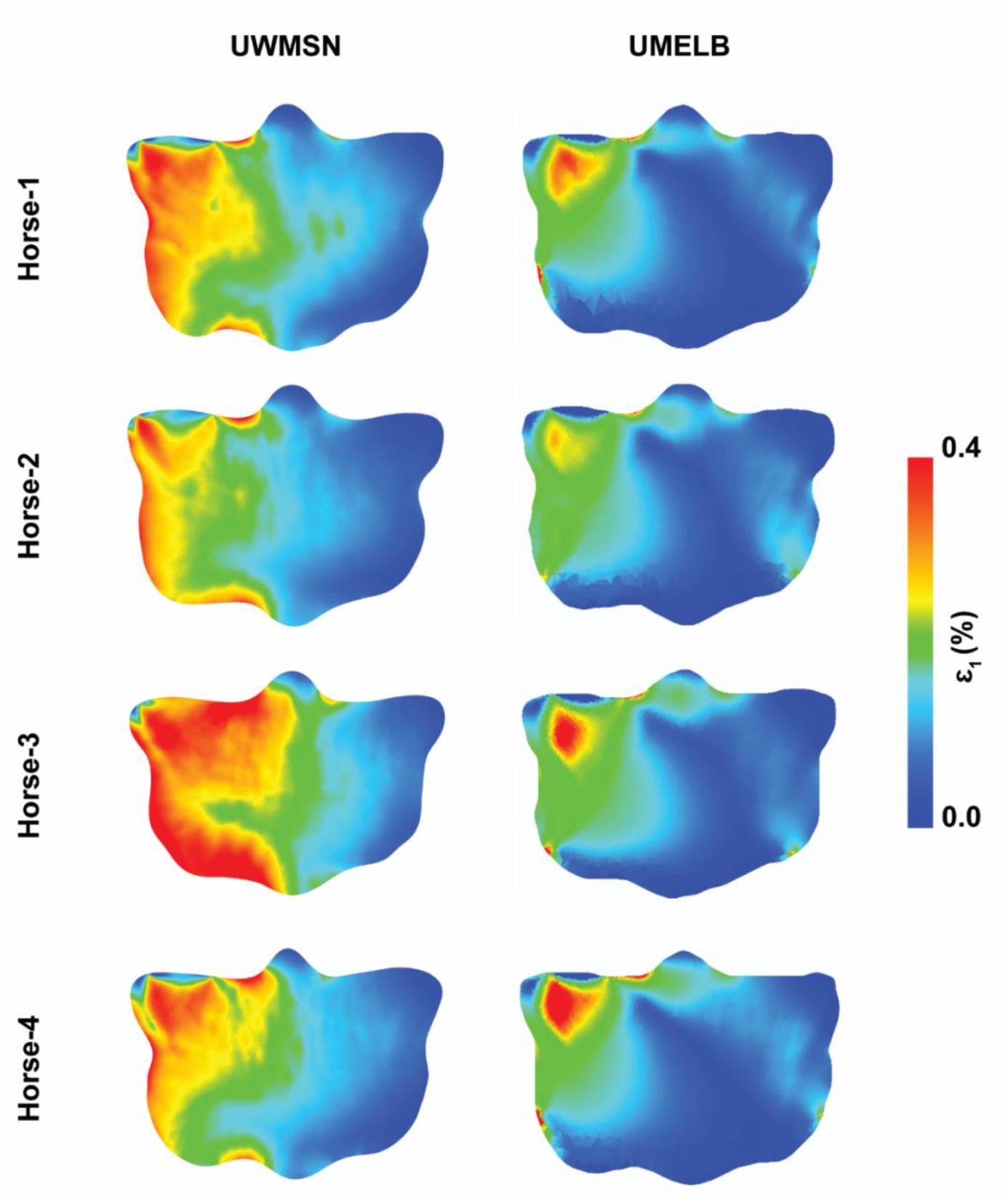
Predicted first principal strain in the middle oblique slice of the four MC3 specimens modeled with the UWMSN (left) and the UMELB (right) pipelines.

Both pipelines predicted a slight increase in the PSG first principal strain from the dorsal to the palmar slice (**Figure 3** and **Figure S2**). On average across the four specimens modeled, UMELB predicted smaller PSG strain in all three slices than predicted by UWMSN. The UMELB pipeline resulted in more comparable values to the UWMSN pipeline when using the density-modulus relationship from Martig *et al*. [18] (equation (5), Melb. Mat. in **Figure 3**) due to the lower Young’s moduli assigned by this relationship compared with the relationship from Moshage *et al*. [17], equation (4) (**Figure S1**). The difference in the predicted PSG strain between the two pipelines, using the same material model, was not consistent on a specimen-to-specimen basis. In Horse-3 for example, UWMSN predicted much larger PSG strain than UMELB, but the predictions were much more similar for Horse-2 (**Figure S2**).

**Figure 3.**
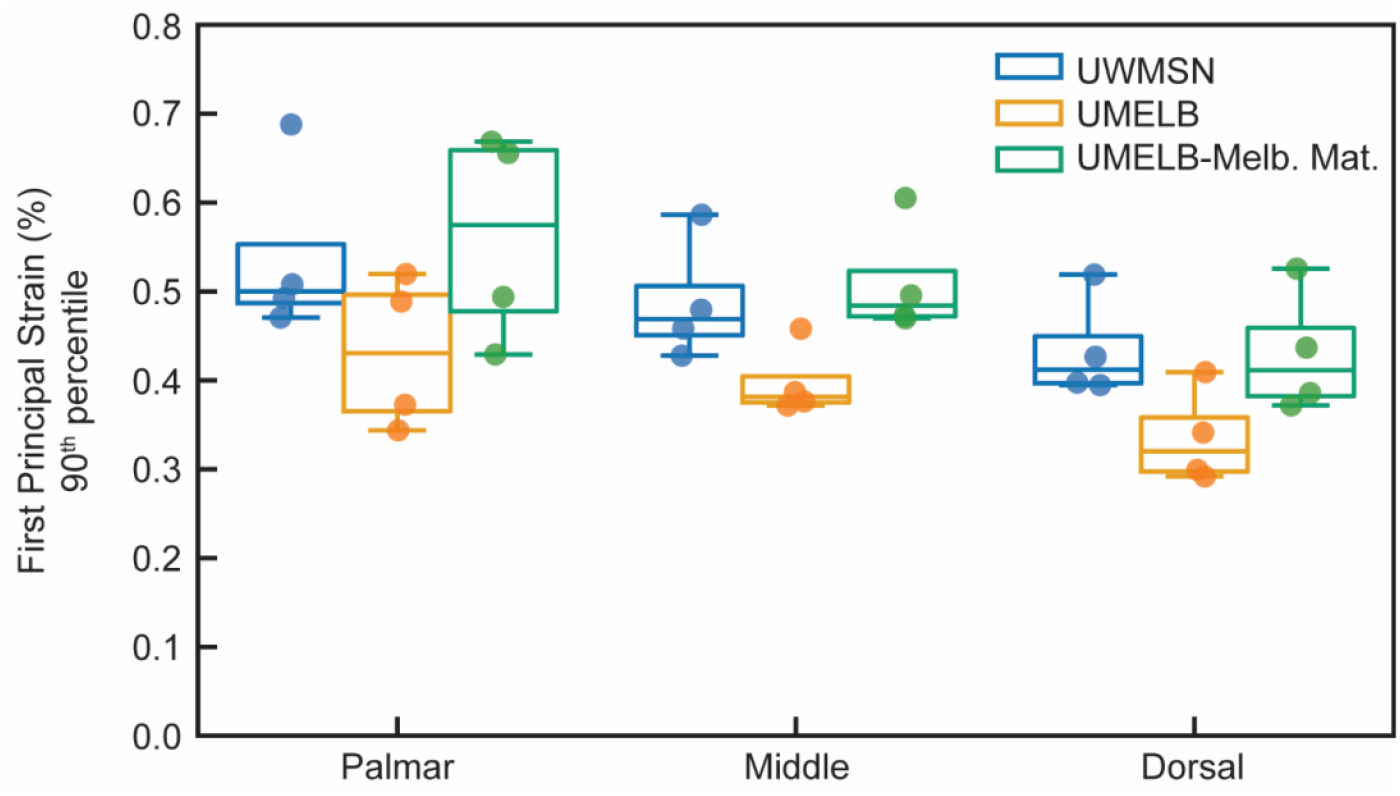
90^th^ percentile first principal strain in the parasagittal groove of the subchondral bone at the three slice locations (palmar, middle, and dorsal), predicted by the UWMSN and the UMELB pipelines for the four third metacarpal specimens modeled. “Melb. Mat.” refers to the density-modulus relationship developed by the Melbourne team, equation (5).

## 4. Discussion

Using the same density-elastic modulus, the slice-based modeling pipeline, UMELB, predicted smaller first principal strain in the PSG, likely due to more limited modes of deformation compared to the full UWMSN 3D FE model, resulting in a stiffer mechanical behavior. Another source of difference between the two modeling pipelines is that UMELB captures the heterogeneity of the bone mineral density in the SCB only in individual slices. Different horses can have subchondral sclerosis with different shapes that would only be captured in a full 3D model, or by modeling each slice, and this difference can create bias in the UMELB pipeline if the subchondral sclerosis has a rare shape. Despite these differences, the predicted strain by the two pipelines remains small, as none of the four specimens used in this study have a large degree of PSG fatigue damage [7], which is a limitation of this study. These specimens, except for Horse-3, had smaller PSG strain under mechanical loading compared to those with high PSG fatigue damage [21].

Experimental validation of the strain predictions enables clinical application of these models and ultimately objective identification of horses at heightened risk of condylar stress fracture. In horses with high degrees of PSG fatigue damage detectable by sCT in the form of subchondral osteolysis [7], prior tuning and validation demonstrated that the UWMSN pipeline requires modification to properly identify the elevation of PSG strain under loading by defining a reduction in the Young’s modulus in the PSG osteolytic region [10]. In addition to this adjustment for horses with high PSG fatigue damage, UWMSN FE models of horses with large subchondral sclerosis required a reduction in Young’s modulus, expected due to the larger degrees of microdamage in such bones [22] that does not affect the bone density and is likely the reason for large variation in the moduli in bone specimens with higher density in the study of Martig *et al*. [18]. Such a reduction of the Young’s modulus in sclerotic bones aligns well with the smaller values reported by Martig *et al*. [18] compared to Moshage *et al*. [17], since the former specifically tested distal MC3 SCB of Thoroughbred racehorses with some degrees of microdamage. This additional validation and tuning has not yet been completed for the UMELB models.

To use these FEA pipelines as a screening injury prevention diagnostic tool, they must be able to identify the elevated PSG strain in horses with PSG fatigue damage, rather than taking the absolute strain value as a predicting measure, especially in the present study where the slice-based specimens are expected to behave in a stiffer manner. We have shown that the UWMSN pipeline with proper tuning of the Young’s modulus in the subchondral sclerotic and lytic bone can predict experimentally measured PSG strain [10] and can classify horses with heightened clinical risk of MC3 condylar stress fracture with high accuracy [23]. It is expected that the UMELB pipeline may require similar tuning to accurately predict strain in a slice-based mechanical testing setup.

## 5. Conclusion

In conclusion, the slice-based modeling pipeline (UMELB) requires less model preparation time and is cheaper to run compared to the full 3D FE model (UWMSN) and once tuned and validated has the potential to identify elevated distal MC3 subchondral strain and ultimately horses with heightened risk of condylar stress fracture.

## Acknowledgements

The authors would like to thank Dr. David Ergun for his help with acquiring the sCT image sets.

## Author Contributions

**Soroush Irandoust:** Conceptualization, Methodology, Software, Formal analysis, Investigation, Writing -Original Draft, Writing -Review & Editing, Visualization. **Fatemeh Malekipour:** Conceptualization, Methodology, Software, Formal analysis, Investigation, Writing -Original Draft, Writing -Review & Editing, Visualization. **R. Christopher Whitton:** Conceptualization, Writing -Review & Editing, Funding acquisition. **Peter Muir:** Conceptualization, Methodology, Formal analysis, Investigation, Resources, Writing -Original Draft, Writing -Review & Editing, Supervision, Project administration, Funding acquisition. **Peter Vee-sin Lee:** Conceptualization, Methodology, Formal analysis, Investigation, Resources, Writing -Original Draft, Writing - Review & Editing, Supervision, Funding acquisition. **Corinne R. Henak:** Conceptualization, Methodology, Formal analysis, Investigation, Resources, Writing -Original Draft, Writing - Review & Editing, Supervision, Funding acquisition.

