## Supplementary Document for "Comparison of two different finite element modeling pipelines for virtual mechanical testing of the distal third metacarpal bone in Thoroughbred racehorses"

Manuscript for *Journal of Biomechanical Engineering*

2025

Soroush Irandoust[1,2,#], Fatemeh Malekipour[3,#], R. Christopher Whitton[4*], Peter Muir[1*], Peter Vee-Sin Lee[3*], Corinne R. Henak[2,5,6*]

[1]Department of Surgical Sciences, School of Veterinary Medicine, University of Wisconsin-Madison, Madison, WI 53706, USA
[2]Department of Mechanical Engineering, University of Wisconsin-Madison, Madison, WI 53706, USA
[3]Department of Biomedical Engineering, University of Melbourne, Parkville, VIC, 3010, Australia
[4]Equine Centre, Melbourne Veterinary School, University of Melbourne, Werribee, VIC, 3030, Australia
[5]Department of Biomedical Engineering, University of Wisconsin-Madison, Madison, WI 53706, USA
[6]Department of Orthopedics and Rehabilitation, University of Wisconsin-Madison, Madison, WI 53705, USA

[#]These authors contributed equally to this work.

**Keywords:** Thoroughbred racehorses, third metacarpal bone, condylar stress fracture, finite element analysis

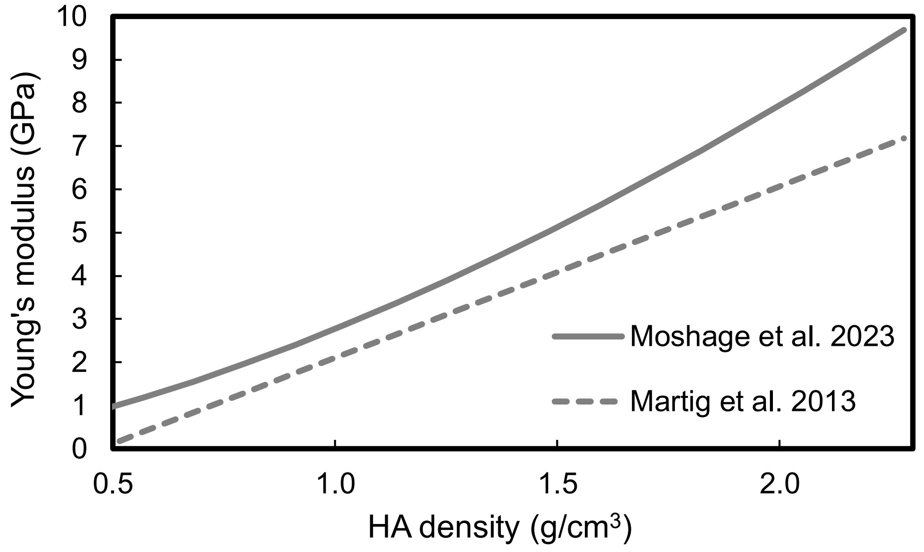


**Figure S1.** Comparison of the two density-modulus relationships used by the UMELB pipeline. UWMSN used the relationship from Moshage *et al.* 2023 only.

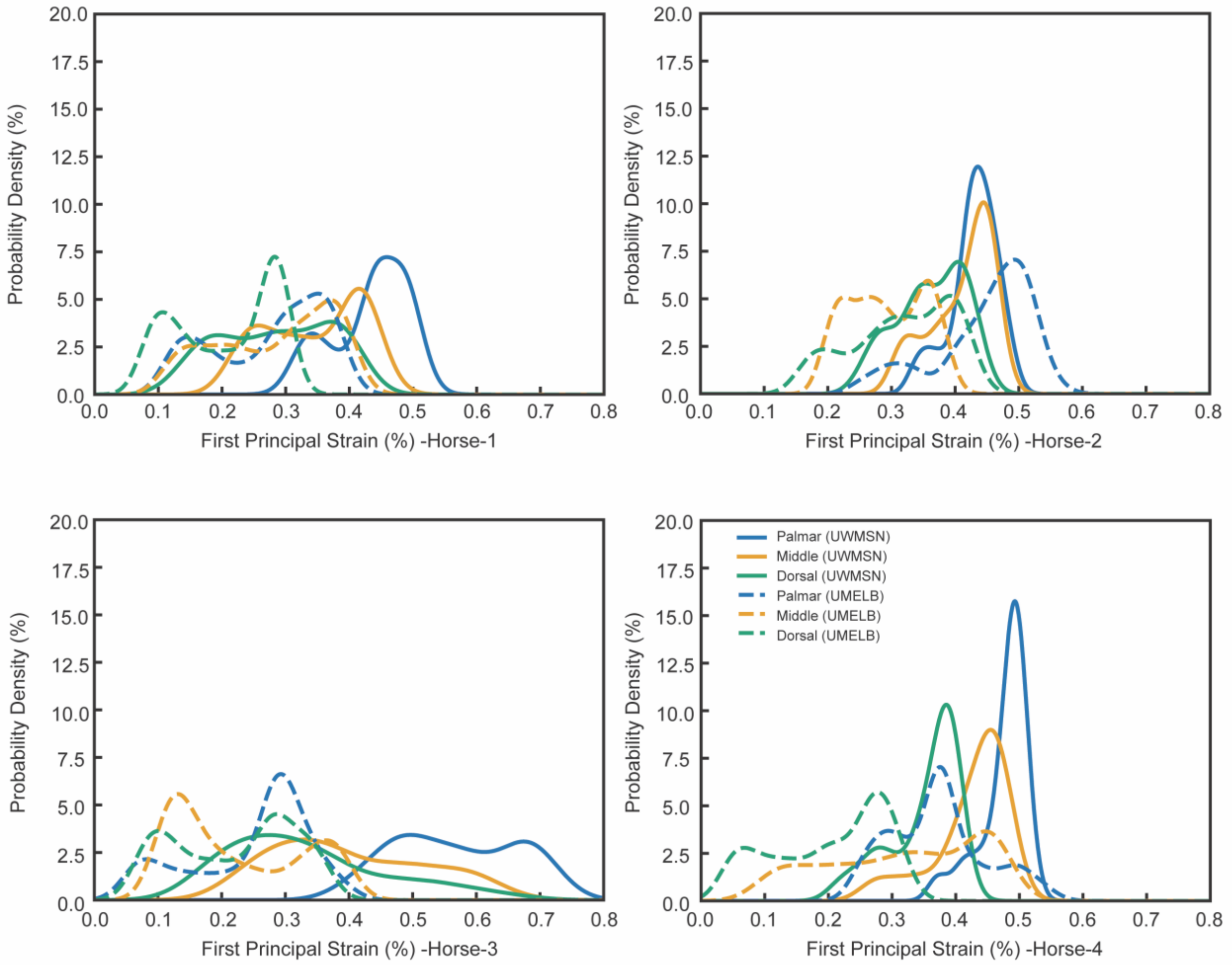


**Figure S2.** Histograms of the UWMSN and UMELB predicted PSG first principal strain in the palmar, middle, and dorsal slices of the four specimens modeled.
